# The geometry of cooperation: decoding microbial interactions

**DOI:** 10.64898/2025.12.22.696080

**Authors:** Stefan Müller, Michael Predl, Diana Széliová, Jürgen Zanghellini

## Abstract

Bridging the gap between mechanistic models of metabolism and ecological theory remains a key challenge in understanding interactions within microbial communities. We propose a geometric framework for analyzing community metabolism, based on constraint-based modeling. By extending pathway analysis methods from single organisms to multi-species systems, we define the *community metabolic space* as the set of all feasible fluxes between species and their environment, conditional on growth rate and medium composition. Embedded within a nonlinear geometry, this space forms a polytope whose vertices represent the minimal building blocks of community metabolism from which every feasible solution can be constructed. Strikingly, these *elementary community flux modes* invite direct ecological interpretation – as specialist, commensalist, or mutualist modes of cooperation. Furthermore, we find that mutualism is either *isotypic* (arising from a minimal mutualistic behavior) or *anisotypic* (emerging from a combination of reciprocal commensalists). This distinction demonstrates that bidirectional cross-feeding alone is insufficient to determine the ecological interaction type. Our framework also offers significant potential for applications. Because it does not rely on optimization, powerful unbiased tools from metabolic engineering, such as production envelopes and minimal cut sets, may be extended to microbial communities. Taken together, this perspective aims to unify ecological and metabolic viewpoints by linking interaction types to the geometric structure of the community metabolic space, thereby laying the foundation for a deeper understanding of community structure, function, and design.

---

Microbial communities thrive by exchanging metabolites and coordinating metabolic tasks, forming complex networks that underpin ecosystem function (1, 2), host physiology (3, 4), and biotechnological processes (5–7). Yet, the principles that govern microbial interactions remain poorly understood and difficult to decipher. Although omics technologies can identify community members and their metabolic capabilities, they offer little insight into the physicochemical conditions that shape microbial interactions (8, 9).

Here, we argue that constraint-based metabolic modeling (CBM) of microbial communities enables a geometric interpretation that reveals the full range of interaction patterns emerging under metabolic constraints. This framework allows us to characterize key ecological relationships – such as mutualism, commensalism, and specialization – as geometric features of the metabolic solution space, offering a new lens on the architecture of microbial cooperation.

## Ecological interactions from metabolic exchange

A common approach to modeling microbial interactions treats them as signed, pairwise relationships between species, describing how the abundance of one species influences the growth of another. In this framework, positive values indicate beneficial effects, negative values represent inhibition or competition, and zero denotes neutrality. These simplified abstractions underlie a broad range of ecological models, including generalized Lotka-Volterra equations, interaction networks, and agent-based simulations, where species abundances change over time based on either fixed or dynamic interactions (10–14).

While such models have provided valuable insights into community dynamics, including stability (15, 16), coexistence, and resilience (17, 18), they remain fundamentally phenomenological: interactions are imposed as external inputs rather than emerging from the underlying metabolic processes. As a result, these models do not explain how specific interaction patterns arise from the metabolic capabilities and constraints of the community members themselves.

## A Geometric View of Cellular Metabolism

To move beyond phenomenological models and towards a mechanistic framework for understanding microbial interactions, CBM offers a powerful approach. For single organisms, CBM has become a cornerstone of systems-level metabolic modeling (19– 23). Its strength lies in our ability to reconstruct metabolism directly from genomic data and to encode the biochemical reaction network of an organism (say *i*) in a stoichiometric matrix ***N*** ^*i*^ (24). This matrix links metabolites to reactions and defines the structure of the metabolic network. Each column quantifies the net production or consumption of metabolites in a reaction. For convenience, we order the columns such that the last represents biomass formation, accounting for all precursor molecules required for growth, including macromolecules such as enzymes, DNA, and RNA.

An organism’s metabolic state is captured by the flux distribution ***v***^*i*^, the vector of net reaction rates, and the biomass flux, which equals the cellular growth rate *µ*_*i*_ scaled by 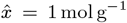 to ensure correct units. By imposing physicochemical constraints, including the steady-state assumption,

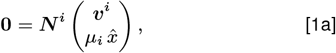

and flux bounds (for example, reflecting reaction irreversibility),

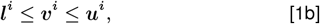

the model restricts fluxes to biologically feasible states, all without requiring detailed kinetic parameters.

### Box 1.

**Polyhedral geometry**

A convex *polyhedron* arises from the intersection of (finitely many) half-spaces, expressed as inhomogeneous linear inequalities. One inequality (for the variable ***x***) can be written as ***a***^*T*^ ***x*** ≥*b*, and hence a polyhedron can be defined by the system

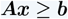

with a matrix ***A*** and a vector ***b*** (the inhomogeneity). A bounded polyhedron is called a *polytope*. A *polyhedral cone* arises from homogeneous linear inequalities and can be written as ***Ax*** ≥**0**. A cone is pointed if it does not contain any line (through the origin).

Alternatively, a polytope is given by convex sums of finitely many vertices, and a pointed polyhedral cone is given by sums of finitely many extreme rays (or, equivalently, by nonnegative sums of representative vectors on the extreme rays).

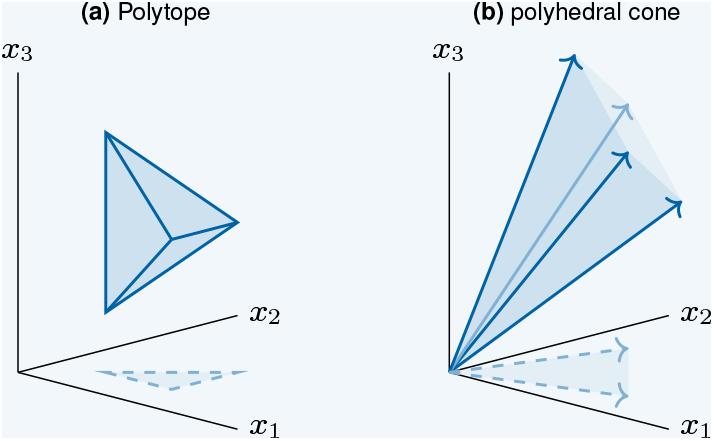

The projection of a polytope (to a subset of its coordinates) is a again a polytope, and the projection of a polyhedral cone is again a polyhedral cone. As simple examples, we show a polytope and a polyhedral cone in three dimensions and their projections to two dimensions in Panel (a) and (b), respectively.

The solution space defined by Eq. [1] is known as the flux polyhedron, whereas Eq. [1a] (together with irreversibility constraints) defines the famous flux cone (25). This space represents all steadystate flux distributions that satisfy the cell’s metabolic capabilities. Fig. 1b shows a projection of this space for a simple cell growing on two substrates, illustrating how substrate uptake shapes the achievable growth rate. For a primer in polyhedral geometry, see Box 1.

**Fig. 1.**
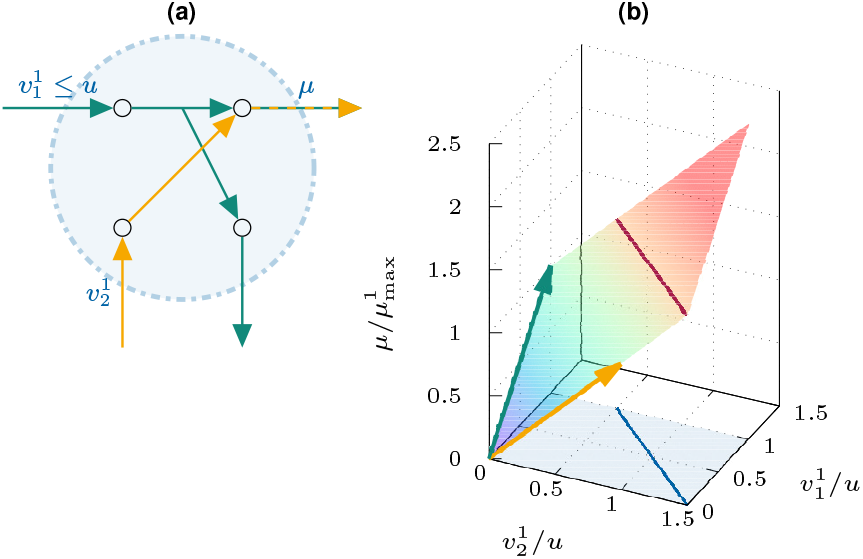
A simple metabolic model of a single microbial species^*‡*^ : (a) The cell takes up a primary substrate from the medium at a rate 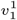, with an upper bound 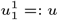 converts this substrate into a biomass precursor and a by-product and excretes the latter. It can also take up a secondary substrate at a rate 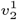 (without an explicit upper bound) to produce biomass. (b) The flux polyhedron describes the set of feasible fluxes for substrate uptake and biomass production. It is defined by two EFVs, shown in green and orange; the corresponding pathways are indicated in Panel (a). Fluxes are scaled by the maximum uptake rate of the primary substrate, *u*, and growth rates are scaled by the maximum growth rate on the primary substrate, 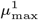. The red line segment indicates a fixed growth rate of 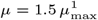 Projections to the exchange fluxes 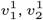 are shown in blue. ^‡^Here, “species” refers to a microbial population sharing a uniform metabolic repertoire.

Beyond visualization, this example reveals a fundamental geometric property: after intersection with an orthant, a region with fixed flux directions, the extreme rays of the flux cone or the vertices of the flux polyhedron correspond to minimal functional metabolic units known as elementary flux modes (EFMs) or elementary flux vectors (EFVs), respectively. Each EFM or EFV represents a non-decomposable steady-state pathway that carries out distinct metabolic functions (26–28), like biomass synthesis or growth-coupled product formation as illustrated in Fig. 1. Moreover, these minimal pathways act as metabolic building blocks; any feasible flux distribution can be composed as a nonnegative or convex linear combination of them. Crucially, if a cell lacks a particular EFM, it cannot perform the associated metabolic function, regardless of how other fluxes are adjusted. For example, the cell in Fig. 1 cannot produce the by-product alone (without growing) because it lacks a corresponding EFM.

## A Geometric View of *Community* Metabolism

Building on this successful foundation, CBM extends naturally to microbial communities (29–31), where metabolic interactions emerge from coupling individual metabolic networks under shared environmental and physicochemical constraints.

Yet, combining multiple metabolic models and rescaling their fluxes introduces a fundamental challenge: as detailed below, the problem becomes nonlinear (32, 33). Unlike the neat flux cone of single organisms, community-level models lose this simple geometric structure. The resulting solution space becomes more complex and computationally demanding, and has so far defied straightforward biological interpretation.

Despite this complexity, the question arises whether we can still map the space of feasible interactions. For example, can we identify mutualism, commensalism, or specialization not as “phenomenological” ecological behaviors, but as intrinsic “elementary modes” within a rigorous mathematical structure? Such an understanding can bring clarity to the diverse ecological interactions within microbial communities.

To explore microbial interactions within a CBM framework, one key assumption must hold: the entire system must be at steady state.

1. For an individual species, this means that intracellular metabolite concentrations remain constant over time, cf. Eq. [1a].
2. Within a community, the steady-state condition also extends to the relative abundance of each species. *Balanced growth*, where all species grow at the same rate, is the only condition under which this is possible. If growth rates differ, relative abundances change, altering the community’s metabolic landscape and breaking the steady-state assumption. Much like in a eukaryotic cell, where organelles such as mitochondria must grow in sync with the rest of the cell to maintain internal balance, microbial communities require coordinated growth to preserve a stable metabolic environment.
3. Steady state also applies to the shared environment, the metabolite concentrations in the culture medium.

Before we formulate the community model mathematically, it is useful to normalize fluxes by the total community biomass, *m* = ∑_*i*_ *m*_*i*_, rather than by the individual biomasses *m*_*i*_. This scaling places all species’ fluxes on a common reference frame, enabling their contributions to be expressed within a single, unified stoichiometric system. Each species’ intracellular flux vector ***v***^*i*^ is weighted by its biomass fraction, *γ*_*i*_ = *m*_*i*_*/m*, where ∑_*i*_ *γ*_*i*_ = 1. This yields the community-scale fluxes 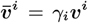 which reflect each species’ relative contribution to the overall community metabolism.

Now, we can formally write the steady-state assumption for species abundances as *γ*_*i*_ = *constant* and the resulting balanced growth condition as *µ*_*i*_ = *µ* for all *i*, that is, all growth rates are equal. Further, we can write the steady-state assumption for the shared environment as

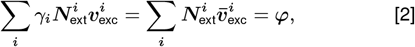

where the additional stoichiometric matrix 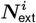 links metabolites in the shared environment (the medium) to exchange reactions, and the vector 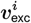 (a subvector of ***v***^*i*^) denotes the corresponding exchange fluxes. The vector **φ** accounts for outflows (*φ*_*j*_ ≥0) from and inflows (*φ*_*j*_ ≤ 0) to the shared environment. This constraint ensures that the net production and consumption of metabolites in the medium by the community matches the environmental inflows and outflows and hence guarantees steady state.

For simplicity, we model the environment as an unlimited reservoir: while the set of externally supplied substrates is specified, the community-level consumption of any substrate *j* is unconstrained (*φ*_*j*_ ≤ 0). Of course, not every species grows on every substrate, and substrate uptake is limited by each cell’s maximum capacity. Similarly, the total excretion of any product *j* from the shared environment into the reservoir is unrestricted (*φ*_*j*_ ≥0). This setup allows us to constrain the directions of exchange with the reservoir, but not their magnitudes; that is, it ensures that environmental availability does not limit the community’s metabolic capacity. Instead, it is the physiology of individual species that sets limits, for example, upper bounds for substrate uptake.

After multiplication with *γ*_*i*_, we combine the intracellular steadystate constraints, Eq. [1], with the extracellular metabolite balance, Eq. [2], thereby using fluxes on community level and assuming an unlimited reservoir of specific substrates. Altogether, we obtain a concise set of equations and inequalities defining the feasible metabolic states of a microbial community under balanced growth.

Equation [3]. Community model under balanced growth

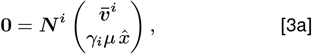

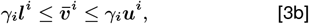

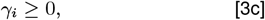

for *i* = 1, …, # species, and

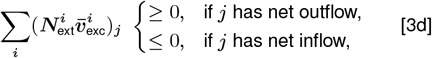

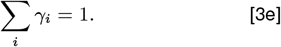

Extending single-species models, the community model is formulated in terms of the community growth rate *µ*, the biomass fractions *γ*_*i*_, and the scaled flux vectors 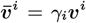 In (34), we further eliminate internal fluxes, that is, we project the model to exchange fluxes (and abundances). Thereby, we reduce dimensions, in particular, we only consider the variables that represent microbial interactions. For conceptual simplicity, we omit this step here.

The community solution space is inherently non-linear and non-polyhedral and hence more complex than the flux cones and polyhedra of individual species. This complexity arises from the bilinear coupling between the biomass fractions *γ*_*i*_ and the community growth rate *µ* in Eq. [3a]. However, for any fixed value of *µ*, the model becomes linear in *γ*_*i*_ and 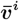 Alternatively, fixing the biomass fractions *γ*_*i*_ yields a linear system for *µ* and 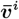

### Example

To visualize this structure, we consider a community of two simple species (see Fig. 2a). The cells are “structurally identical”, but differ in their exchange with the environment. For simplicity, we assume equal maximum substrate uptake rates, 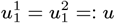 leading to equal individual maximum growth rates, 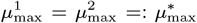 Fig. 2b shows a projection of the highdimensional space of the feasible metabolic states onto three variables: the community growth rate *µ*, the relative abundance *γ*_1_ of species 1, and its substrate uptake rate 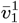. Each horizontal slice corresponds to a fixed *µ*, and defines a convex polytope of feasible states. As *µ* increases, these polytopes change: from a line segment at *µ* = 0, through triangles, to a single point at the maximum growth rate 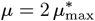 Stacking these slices across all *µ* produces a non-convex, non-polyhedral geometry. This shape highlights the community’s metabolic flexibility at intermediate growth rates and the tight constraints at the growth extremes. The doubling of the maximum community growth rate, relative to that of individual species, reflects the beneficial interactions between cells which enable the collective to achieve higher performance than any species could independently.

**Fig. 2.**
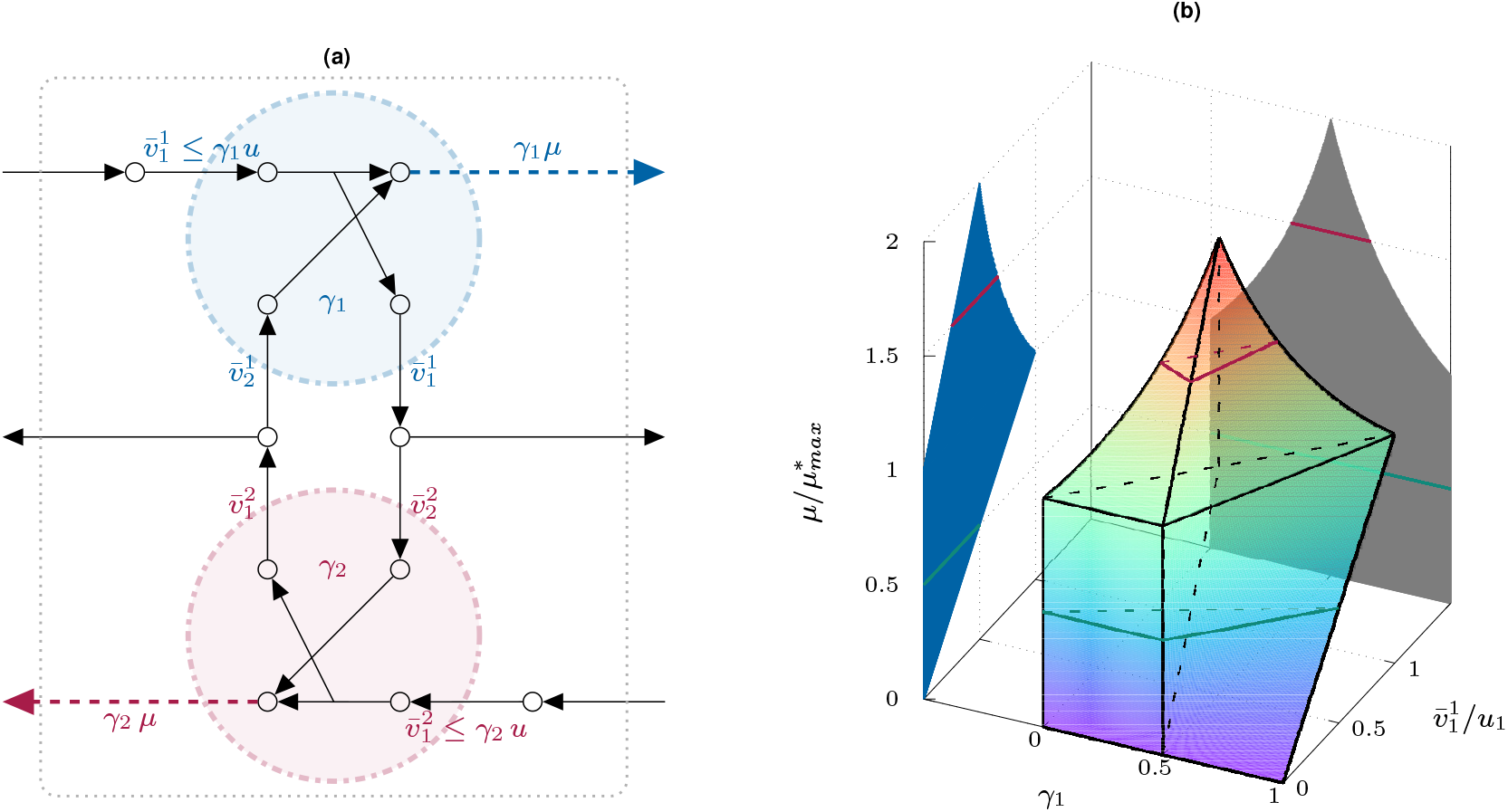
Community of two simple microbial species: (a) The first cell (blue) takes up a substrate from the medium (subject to an upper bound), converts it into biomass precursor and a by-product and secretes the latter. Alternatively, it can grow on another substrate, available only if secreted by the second species. The second cell (red) is a “mirror image” of the first, with analogous metabolic capabilities, but different substrates and products. (b) The high-dimensional space of the feasible metabolic states of the community is projected onto three key variables: the growth rate *µ*, the substrate uptake rate of the first species 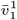, and its relative abundance *γ*_1_. Although the overall geometry is nonlinear in *µ* (and *γ*_1_), fixing *µ* (or *γ*_1_) yields a convex polytope, illustrated here by the green and red triangles at 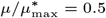 and 1.5 (as well as by the black parallelogram at *γ* = 0.5). For a stepwise construction of the red triangle at 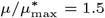, see Box 2.

Fig. 2b also illustrates that not only slices at fixed growth rate *µ* yield convex polytopes, but also those at fixed abundances *γ*_*i*_. Both cross-sections define feasible metabolic states of the community. The two families of slices reflect the bilinear structure of the underlying community model, Eq. [3a] (linear in *µ* for fixed *γ*_*i*_, and linear in *γ*_*i*_ for fixed *µ*), embedded within a globally nonlinear geometry.

In order to better understand how the community space in Fig. 2b arises, we outline a step-by-step construction for a fixed growth rate 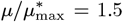 in Box 2. As described by Eqs. [3a], [3b], and [3c], each species contributes its phenotypic metabolic capabilities, scaled continuously across all possible relative abundances. Geometrically, this scaling defines convex cones of feasible metabolic states, bounded further by the constraint *γ*_*i*_ ≤1. The coupling of exchange fluxes in Eq. [3d] determines how these individual spaces align and overlap to support a feasible community. The resulting intersection forms the final community solution space.

## Analyzing community metabolism

With the structure of the community metabolic space under balanced growth established, we turn to methods for analyzing it. Most approaches, analogous to classical flux-balance analysis (FBA) (35), focus on identifying states that maximize the growth rate *µ*. One of the earliest, community flux-balance analysis (cFBA) (33), treats biomass fractions as parameters. For each parameter set, it solves a linear program to maximize *µ*, effectively scanning the solution space along slices for fixed *γ*_*i*_. By contrast, for each growth rate, SteadyCom (32) solves for the corresponding biomass compositions *γ*_*i*_, exploring the space along slices for fixed *µ*. This approach avoids the high-dimensional scan over biomass fractions and scales better to large communities.

The optimal growth rate can often be achieved by multiple distinct configurations. To resolve this ambiguity, methods like OptDeg (36) introduce a secondary objective that selects metabolic states where each species maximizes its own biomass yield. This “egoistic” refinement narrows down the solution space to configurations where no species sacrifices growth for the benefit of others, revealing how far individual efficiency can account for community-level outcomes. Yet, all three methods remain confined to the growth-maximizing paradigm, offering little insight into the suboptimal regions of the feasible metabolic state.

### Unbiased approach

In order to move beyond growth-maximizing states and to characterize the full community solution space, we draw on tools from metabolic pathway analysis (26, 27, 37). In single-species models, the feasible steady-state fluxes form a polyhedral cone, the so called flux cone. The extreme rays of this cone, more precisely of its intersection with an orthant, correspond to EFMs, the irreducible metabolic steady-state pathways.

A similar structure emerges in the community setting. For fixed growth rate *µ*, the community model Eq. [3] can be written as a homogeneous linear system, with all variables on the left-hand side and zero on the right-hand side. This defines a (community) cone in the variables *γ*_*i*_ and 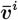, analogous to the classical flux cone for a single organism. However, unlike in single-species models, where flux bounds typically convert the flux cone into a polytope, here the bounds are absorbed into the cone structure. The only inhomogeneous constraint Eq. [3e], which requires biomass fractions to sum to one, describes a transversal intersection of the cone which yields the polytope of feasible community states.

The extreme rays of the community cone represent minimal community-level flux distributions, which we term elementary community flux modes (ECFMs). Unlike EFMs, ECFMs depend on the growth rate *µ*. Yet, just as EFMs reveal the architecture of cellular metabolism, ECFMs capture the fundamental metabolic interactions that enable different species to collectively grow at the same rate. In fact, any feasible community flux distribution can be expressed as a convex linear combination of these ECFMs which makes them the natural building blocks for analyzing the geometry of community metabolism. Since ECFMs are structurally analogous to EFMs, they can be readily computed using existing software for metabolic pathway analysis (38–40), as demonstrated in the accompanying Jupyter notebook (41).

#### Box 2.

**Constructing the community metabolic space**

We illustrate how metabolic interactions shape the community metabolic space. We consider the community of two simple microbial species in Fig. 2a and construct (a projection of) this space at a fixed community growth rate of 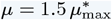. In the end, we obtain a single slice of the full space, namely the red triangle in Fig. 2b.

In Panel (c), we instantiate the community model, Eq. (3), for the exchange fluxes and abundances of species 1 and 2, including equality constraints for cell growth [G], upper bounds for substrate uptake [B], and nonnegativity constraints for fluxes and abundances [N], highlighted in blue for cell 1 and in red for cell 2. The model further involves inequalities for the exchange fluxes [X], since metabolites secreted by one species may only be partially consumed by the other, causing an outflow to the reservoir. Finally, the model includes a convexity constraint for the abundances [C]. For simplicity, we assume [N] and [C] in all other panels and do not state them explicitly.

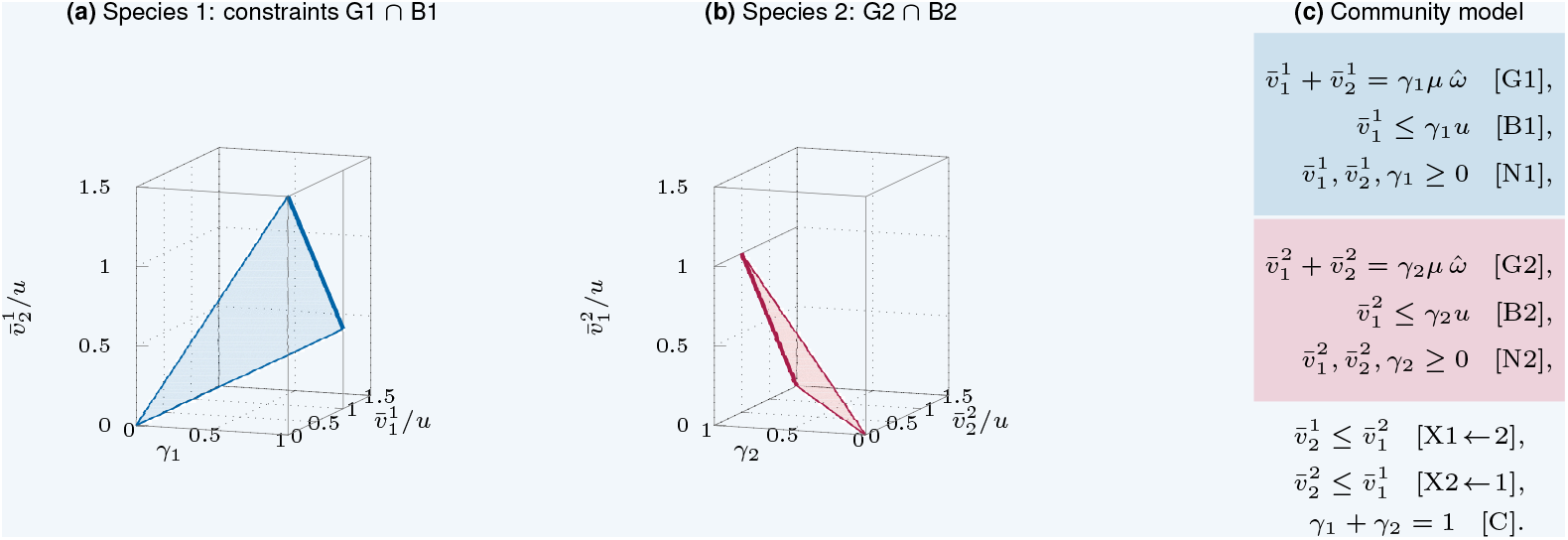

In Panels (a) and (b), we first consider each species individually, formulating constraints in terms of its scaled fluxes and abundance. At the fixed growth rate and full abundance (*γ*_1_ = 1), the feasible metabolic space for species 1 equals the thick blue line segment in Panel (a), cf. Fig. 1b. As the abundance *γ*_1_ varies between 0 and 1, the space scales proportionally. It forms a convex polyhedral cone, indicated by the light blue triangle. The scaling arises since the constraints [G1], [B1], and [N1] are homogeneous in the variables 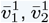, and *γ*_1_ of species 1. The same reasoning applies to species 2, as shown in Panel (b).

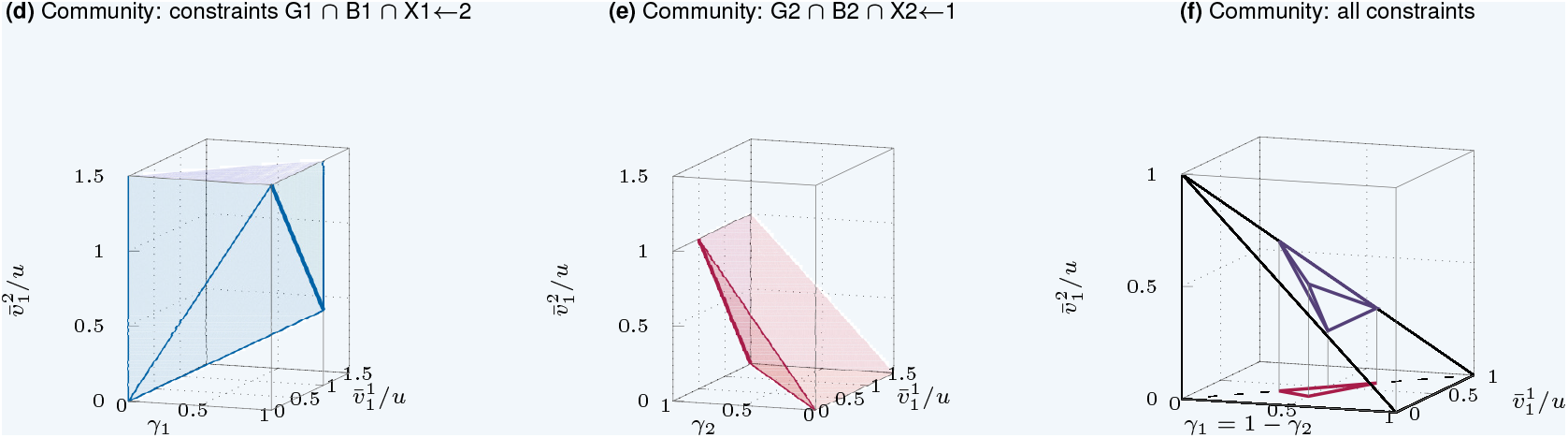

Next, we account for the fact that the alternative substrate of species 1 is produced by species 2, which introduces the “coupling” 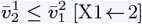. To illustrate this constraint, we have to use variables of both species, and we choose to project to the primary substrate uptake fluxes 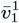 and 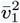 and the abundance *γ*_1_. When going from Panel (a) to Panel (d), we replace the variable 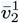 by 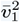, and the feasible metabolic space in Panel (d) equals the prism above the triangle in Panel (a). Note that, we have considered one more constraint, but we also go from individual species to community level, and hence the (projection of the) feasible region has higher dimension. The same applies to species 2 in Panel (e).

The community metabolic space emerges by intersecting the two spaces in Panels (d) and (e) and using the identity *γ*_1_ = 1 − *γ*_2_ [C]. The resulting tetrahedron in Panel (f) captures all feasible metabolic states of the two-species community in Fig. 2a at a fixed growth rate of 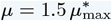. Its projection to the variables *γ*_1_ and 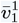 yields the red triangle in Fig. 2b.

### Example (continued)

For the two-species community in Fig. 2a, representative ECFMs for growth rates 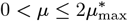 are shown in Fig. 3. For 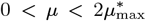, there are always two pairs of ECFMs, each pair representing symmetric community-level flux distributions with the roles of the two species exchanged. (In Fig. 2b, only one ECFM of the pair residing at *γ*_1_ = *γ*_2_ = 0.5 remains a vertex after projection to *γ*_1_ and 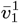, and hence the feasible metabolic states are generated by three points and form a triangle.) At the maximum growth rate, all ECFMs collapse into a single mode, meaning that there is only one possible way for the community to organize itself.

**Fig. 3.**
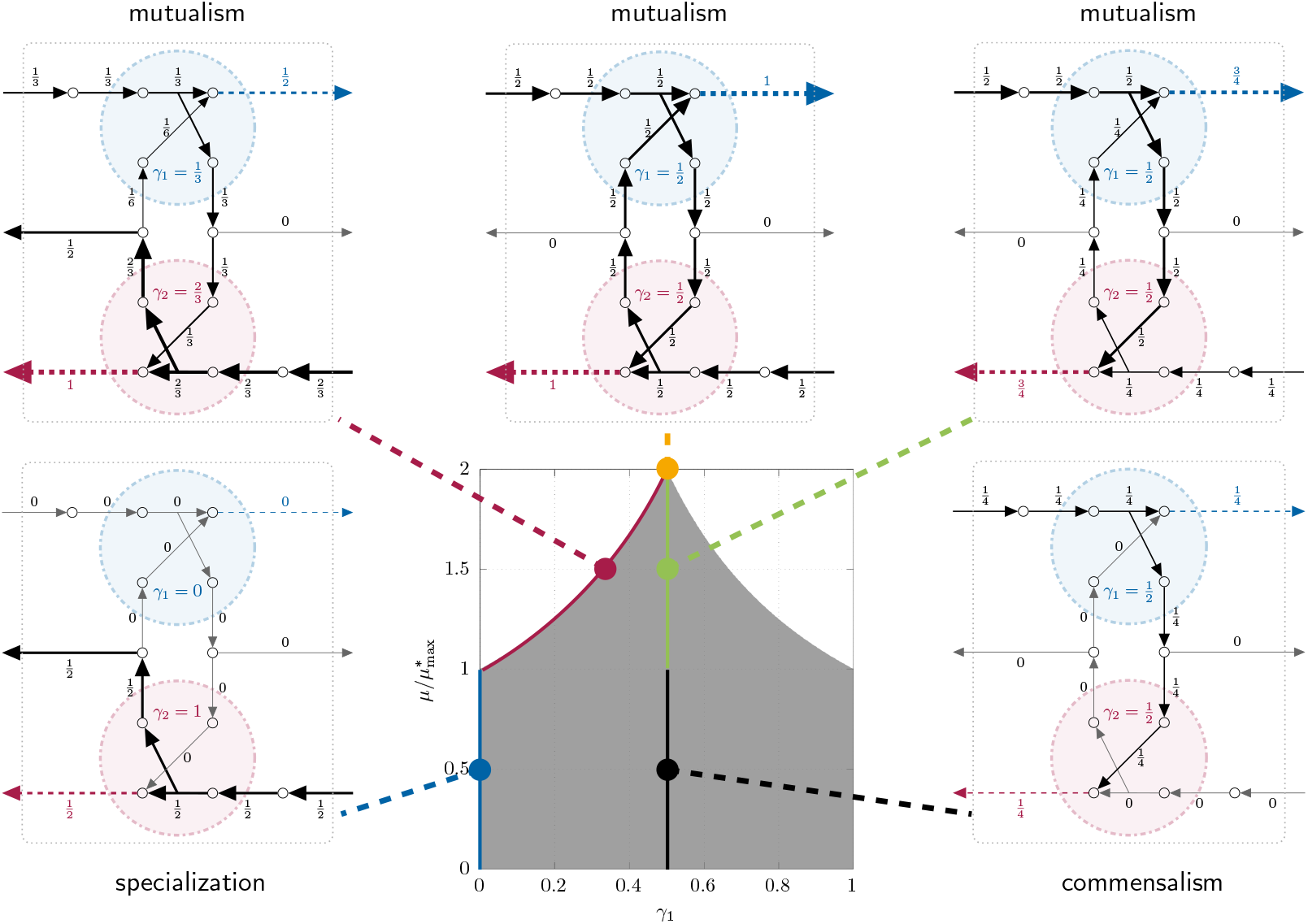
ECFMs and their ecological interpretations for the two-species community in Fig. 2a, shown in the projection of the feasible space to composition and growth rate, cf. Fig. 2b. For each growth rate 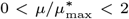 there are four ECFMs; in fact, two pairs of ECFMs that are symmetric regarding the roles of species 1 and 2. (At 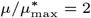, all ECFMs collapse into one strategy.) At 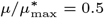, we mark two ECFMs each (by blue, black, red, and green circles), and at maximum growth rate, we mark the unique feasible strategy (by an orange circle). For each ECFM, we show the underlying reaction network with corresponding flux values. Most importantly, the flux patterns of the ECFMs can be ecologically interpreted as mutualism, commensalism, or specialization. The colored continuous lines indicate the position of ECFMs that are qualitatively identical to the representatives with the same color. The areas between ECFMs consist of flux patterns that are linear combinations of ECFMs (with the same growth rate).

Although the flux values in each ECFM vary with the community growth rate *µ*, their qualitative properties remain unchanged within the intervals 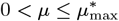 and 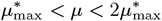, respectively. The set of active constraints and the corresponding flux pattern stays the same throughout each interval. It is therefore sufficient to examine representative ECFMs at 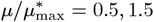, and 2 to capture the distinct modes of community interaction, see Fig. 3.

In the low-growth regime 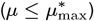, the two pairs of ECFMs reveal two minimal community behaviors. In the first, only one species is present: it consumes substrate from the environment, converts it into biomass, and releases a by-product (which flows out to the reservoir). In the second, both species coexist in a cross-feeding relationship; one takes up a substrate and produces a by-product, while the other relies solely on that by-product for growth. The second behavior is possible only when the species have equal abundance, *γ*_1_ = *γ*_2_. Importantly, within this growth range, the two pairs of ECFMs suffice to generate every other feasible metabolic state of the community as a convex sum.

In the high-growth regime 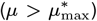, both species take up their substrates, which flow in from the reservoir, and convert them into biomass and by-products. Unlike in the lower regime, each species also takes up the by-product produced by the other, enabling growth beyond 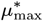, the maximum growth rate of the individual species on the primary substrate. In fact, there are two minimal community behaviors, differing in their metabolic efficiency: in the first, both species take up substrates at maximum capacity, resulting in excess by-product release; in the second, substrate uptake is below maximum and adjusted to prevent by-product outflow. Again, due to symmetry, the second behavior occurs only for *γ*_1_ = *γ*_2_ = 0.5.

### Ecological significance

The ecological interpretation of the minimal community-level flux distributions is striking. In the high growth regime 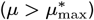, all ECFMs represent mutualism (+/+): each species takes up its primary substrate flowing in from the reservoir and excretes a by-product utilized by the other. This bidirectional cross-feeding allows both species to benefit directly and enables the community to surpass the growth limit of either species alone.

In the low-growth regime 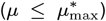, minimal community behaviors include unidirectional cross-feeding, corresponding to commensalism (0/+): one species benefits (+) by consuming the by-product of the other, while the producer remains unaffected (0), as the by-product arises “cost-free” from growth-coupled production. The second possible behavior is specialization, where only one species grows on its primary substrate while the other is not present. These findings show that minimal community states derived by CBM can be directly interpreted as classical ecological interactions (mutualism, commensalism, and specialization). Within this framework, interaction types emerge naturally from the underlying metabolic network structures and environmental constraints, rather than being defined phenomenologically.

### Beyond ecological interaction types: isotypic vs. anisotypic mutualism

Both growth regimes exhibit bidirectional cross-feeding, but with fundamentally different origins, see Fig. 4. In the high-growth regime 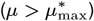, all minimal community-level flux distributions are bidirectional; both species simultaneously produce and consume each other’s by-products (Fig. 4a). Hence, any necessarily bidirectional feasible metabolic state is generated using a bidirectional ECFM, and we call it *isotypic*. In contrast, in the low-growth regime 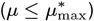, minimal community states involve only unidirectional cross-feeding, and bidirectionality emerges solely as a composition of unidirectional ECFMs (Fig. 4b). Here, bidirectional cross-feeding is *anisotypic*.

**Fig. 4.**
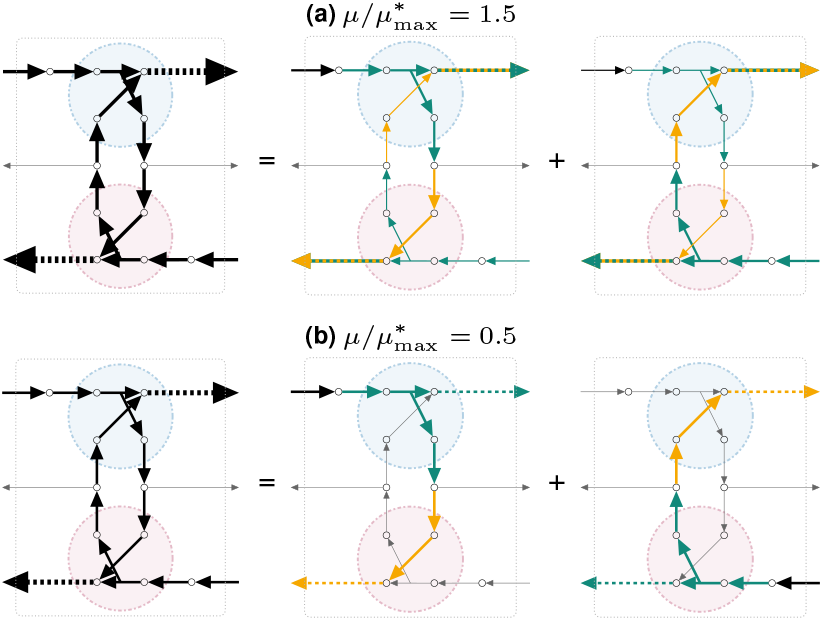
Isotypic (a) and anisotypic (b) decomposition of bidirectional cross-feeding in the two growth regimes of the two-species community in Fig. 2. Green and orange refer to EFMs of individual species.

This highlights that the nature of bidirectional cross-feeding is shaped by community-level constraints. It adds a new dimension to existing classifications of cross-feeding (42), showing that observed bidirectionality alone does not distinguish between isotypic and anisotypic mutualism.

In the example, the distinction is reflected when decomposing ECFMs by species. In the low-growth regime, unidirectional ECFMs decompose into exactly one EFM per species (Fig. 4b); each cell operates via a “pure” minimal metabolic strategy. By contrast, in the high-growth regime, bidirectional ECFMs require each species to coordinate multiple EFMs simultaneously (Fig. 4a), resulting in “mixed” metabolic strategy at the species level.

This observation leads us to hypothesize a potential general principle: commensal ECFMs are decomposable into single EFMs (or EFVs) per species, whereas mutualistic ECFMs may combine several EFMs (or EFVs) at species level.

If true, this would imply a connection between community-level interaction types and the complexity of intracellular metabolic coordination. “Pure” strategies at species level correspond to unidirectional interactions (commensalism), whereas mutualistic strategies may require cells to “mix” multiple metabolic functions. Though speculative, this opens an intriguing direction for future research: to what extent does intracellular metabolic management shape ecological interactions?

## Future Directions

ECFMs offer a powerful lens through which to study metabolic interactions in microbial communities. As minimal, non-decomposable flux modes, they serve as the fundamental building blocks of community metabolism and provide a natural foundation for addressing longstanding questions in microbial modeling and ecology.

One such question concerns the nature of intracellular decision-making: should community members be viewed as egoistic optimizers, or do community-level constraints give rise to flux patterns that resemble altruistic behavior? The optimality degree (OptDeg) has been proposed as a measure of selfishness (36). It quantifies how close the community is to achieving optimal substrate utilization (biomass yield) for all organisms involved. By applying metrics such as OptDeg to ECFMs, one can systematically assess the degree to which individual benefit is sacrificed in each minimal community state.

The ReMIND framework (43) recently introduced a related approach: it reconstructs metabolic interaction networks from individual cellular phenotypes that minimize the number of exchange reactions with the environment. Instead of scanning the full solution space, ReMIND uses integer programming to directly enumerate combinations of these minimal units that optimize substrate utilization, effectively identifying the set with maximum OptDeg.

The emphasis on minimality in methods like ReMIND and ECFM analysis reflects a broader shift toward parsimony as a guiding principle for understanding complex microbial interactions. This focus promotes mechanistic insight and interpretability. Integrating parsimony with economic reasoning can identify likely metabolic exchanges, reveal trade-offs between yield and cooperation, and constrain models to biologically meaningful regions of metabolic space.

Building on this mechanistic foundation, ECFMs also offer a natural interface with game theory (44), as each mode represents a community state, while the ECFMs of individual species serve as discrete strategies. Introducing payoff concepts such as growth rate enables the identification of Nash equilibria within the space of minimal community states, configurations in which no species can unilaterally improve its outcome. While game-theoretic ideas have been applied to metabolic models (45, 46), their integration with the assumption of balanced growth remains largely unexplored. In this context, ECFMs provide a promising framework for investigating the emergence and stability of cooperative behavior.

A further opportunity lies in refining the classification of microbial interactions. Existing approaches are largely phenomenological, relying on network topology or interaction signs (8, 42). In contrast, ECFMs provide a mechanistic perspective that enables decomposition into interpretable building blocks. For example, they make it possible to distinguish between interactions that are additive across organisms and those that are emergent, arising only through collective metabolic activity.

### Broader applications

The ECFM framework also opens new possibilities for metabolic engineering. In particular, methods such as minimal cut sets and production envelopes (47), which have been highly effective for industrial strain design in single-species models, can now be applied to microbial communities. The correspondence between ECFMs and ecological concepts creates opportunities for designing microbial consortia that integrate both intracellular metabolism and interspecies interactions.

Relaxing the assumption of balanced growth across species would substantially broaden the framework’s applicability. For instance, incorporating species-specific death or dilution rates could capture stable communities with heterogeneous growth dynamics. Such extensions would bring the model closer to natural systems, while still preserving the underlying ECFM structure.

While the ECFM approach is primarily focused on microbial communities, it is also applicable to eukaryotes, offering intriguing new modeling opportunities. Through the lens of ECFMs, a eukaryotic cell can be viewed as a community of compartments, allowing one to investigate how their relative abundance shapes overall cellular metabolism.

By placing ECFMs at the core of the framework, we highlight how minimal community behaviors can serve as a unifying scaffold for understanding cooperation in microbial ecosystems and guiding the design of synthetic consortia.

## Computation

ECFMs were computed using efmtool v0.2.1 (40) with input networks in SBML generated via PyCoMo (48). Code for reproducing the small example is available at github.com/diana-sz/community_modes and (41).

## Acknowledgments

This research was funded in whole or in part by the Austrian Science Fund (FWF), 10.55776/P33218, 10.55776/PAT3748324,and 10.55776/COE17, [Cluster of Excellence: Circular Bioengineering](https://www.circularbioengineering.at/).

For open access purposes, the authors have applied a CC BY public copyright license to any author-accepted manuscript version arising from this submission.

